# Influence of learning strategy on response time during complex value-based learning and choice

**DOI:** 10.1101/248336

**Authors:** Shiva Farashahi, Katherine Rowe, Zohra Aslami, M Ida Gobbini, Alireza Soltani

**Affiliations:** Department of Psychological and Brain Sciences, Dartmouth College, NH, United States of America; Dipartimento di Medicina Specialistica, Diagnostica e Sperimentale (DIMES), Medical School, University of Bologna, Bologna, Italy

## Abstract

Measurements of response time (RT) have long been used to infer neural processes underlying various cognitive functions such as working memory, attention, and decision making. However, it is currently unknown if RT is also informative about various stages of value-based choice, particularly how reward values are constructed. To investigate these questions, we analyzed the pattern of RT during a set of multi-dimensional learning and decision-making tasks that can prompt subjects to adopt different learning strategies. In our experiments, subjects could use reward feedback to directly learn reward values associated with possible choice options (object-based learning). Alternatively, they could learn reward values of options’ features (e.g. color, shape) and combine these values to estimate reward values for individual options (feature-based learning). We found that RT was slower when the difference between subjects’ estimates of reward probabilities for the two alternative objects on a given trial was smaller. Moreover, RT was overall faster when the preceding trial was rewarded or when the previously selected object was present. These effects, however, were mediated by an interaction between these factors such that subjects were faster when the previously selected object was present rather than absent but only after unrewarded trials. Finally, RT reflected the learning strategy (i.e. object-based or feature-based approach) adopted by the subject on a trial-by-trial basis, indicating an overall faster construction of reward value and/or value comparison during object-based learning. Altogether, these results demonstrate that the pattern of RT can be informative about how reward values are learned and constructed during complex value-based learning and decision making.

## Introduction

Investigations of response time (RT) have long been the focus of human psychophysics studies in order to distinguish between alternative mechanisms underlying mental processes (Donders, 1868; Luce, 1986; Sternberg, 1969b). Similarly, psychologists and neuroscientists have used the pattern of RT in different experimental conditions to infer neural processes underlying various cognitive functions. These include but are not limited to: visual processes (Breitmeyer, 1975; Duncan & Humphreys, 1989; Tanaka & Shimojo, 1996; Treisman & Sato, 1990; Tynan & Sekuler, 1982), attention (Joseph, Chun, & Nakayama, 1997; Krummenacher, Grubert, & Müller, 2010; Krummenacher, Müller, & Heller, 2001; Treisman & Sato, 1990; Wolfe, Cave, & Franzel, 1989), memory (Sternberg, 1969a; Unsworth & Engle, 2005), and decision making (Logan, Cowan, & Davis, 1984; Pleskac & Busemeyer, 2010; Ratcliff, Van Zandt, & McKoon, 1999; Spiliopoulos & Ortmann, 2016). Here we asked whether RT could also be used to infer how reward values are learned and constructed in the brain.

In naturalistic settings, humans must use information about an object, such as its features (i.e. color, taste, texture), to determine whether it is rewarding. This is because one cannot learn the reward values for a large number of options that exist in the real world. For example, the features of fruit in a school cafeteria line may allow students to choose between many types of fruit available. A green, squishy pear may be compared to a red, crispy apple, based on what one knows about these features. Individual features, however, are not always predictive of reward; whereas a green apple may be tasty, a green banana may not. Therefore, in order for learning to adapt to different situations, it must be adjusted to incorporate reward feedback differently depending on the nature of the environment. Currently, extensive literature addresses how humans exploit various heuristics to tackle real-world decision-making and learning tasks (Fellows, 2006; Gigerenzer & Goldstein, 1996; Gilovich, Griffin, & Kahneman, 2002; Jocham et al., 2016; Niv et al., 2015). We have recently shown that humans use a heuristic strategy to tackle learning reward values in dynamic multi-dimensional environments (Farashahi, Rowe, Aslami, Lee, & Soltani, 2017). More specifically, rather than directly estimating the reward value of individual options based on reward feedback (object-based learning), subjects estimate average reward values of features and then combine these values to estimate reward values for individual options (feature-based learning). Importantly, these learning strategies not only require different representations of reward values and updates based on reward feedback but also involve distinct methods for construction or comparison of reward values to make a decision.

We hypothesized that the way reward values are constructed and compared could influence how quickly a choice between two alternative options can be made. To examine this influence, we considered three stages involved in value-based decision making. First, objects and/or their features need to be detected and identified in order to evoke the corresponding representations of reward values. Second, in the case of feature-based learning, the individual values of relevant features should be combined to construct the final value of an object. Third, the final values of two alternative objects should be compared to make a decision (Lee, Seo, & Jung, 2012; Soltani & Wang, 2006, 2008; Sugrue, Corrado, & Newsome, 2005).

Because the choice during a given trial depends primarily on the difference between the reward values of the two options on that trial (Barraclough, Conroy, & Lee, 2004; Corrado, Sugrue, Seung, & Newsome, 2005; Lee et al., 2012; Louie & Glimcher, 2010; Seo, Barraclough, & Lee, 2007; Sutton & Barto, 1998), decision making should be easier and faster when this difference is larger. This predicts that RT should decrease with the absolute value of the difference in reward values on a given trial. Moreover, if reward values of objects are directly estimated/learned (object-based learning), retrieval of reward values only requires identification of the two objects. In the case of feature-based learning, however, the subject has to construct reward values of objects after retrieving their feature values and combining them. This predicts longer RT when feature-based learning is adopted, indicating that RT could reflect the dominant learning strategy on a trial-by-trial basis when both strategies are present and compete with each other to determine choice.

To test the aforementioned predictions, here we analyzed choice and RT data during a set of multi-dimensional learning and decision-making tasks that can promote different types of learning strategies.

## Materials and methods

### Overview of experiments and approach

Human subjects performed a set of multi-dimensional learning and decision-making tasks. During each trial, the subject selected between pairs of objects (colored shapes or patterned shapes) that would yield reward with different probabilities (Figure 1). These probabilities could stay the same (static environment) or unpredictably change over time (dynamic or volatile environment). Moreover, the relationship between reward probabilities of objects and their features created generalizable or non-generalizable environments. In a generalizable environment, the reward probabilities assigned to different objects could be estimated from their features based on a multiplicative rule. In contrast, in a non-generalizable environment, reward probabilities assigned to different objects could not be accurately estimated using their features. By fitting choice behavior with seven different models, we identified which of the two classes of learning strategies (feature-based or object-based) was adopted by individual subjects during each experiment. We then examined prediction of the best object-based and feature-based models on a trial-by-trial basis in order to study how RT depended on the adopted learning strategy.

**Figure 1.**
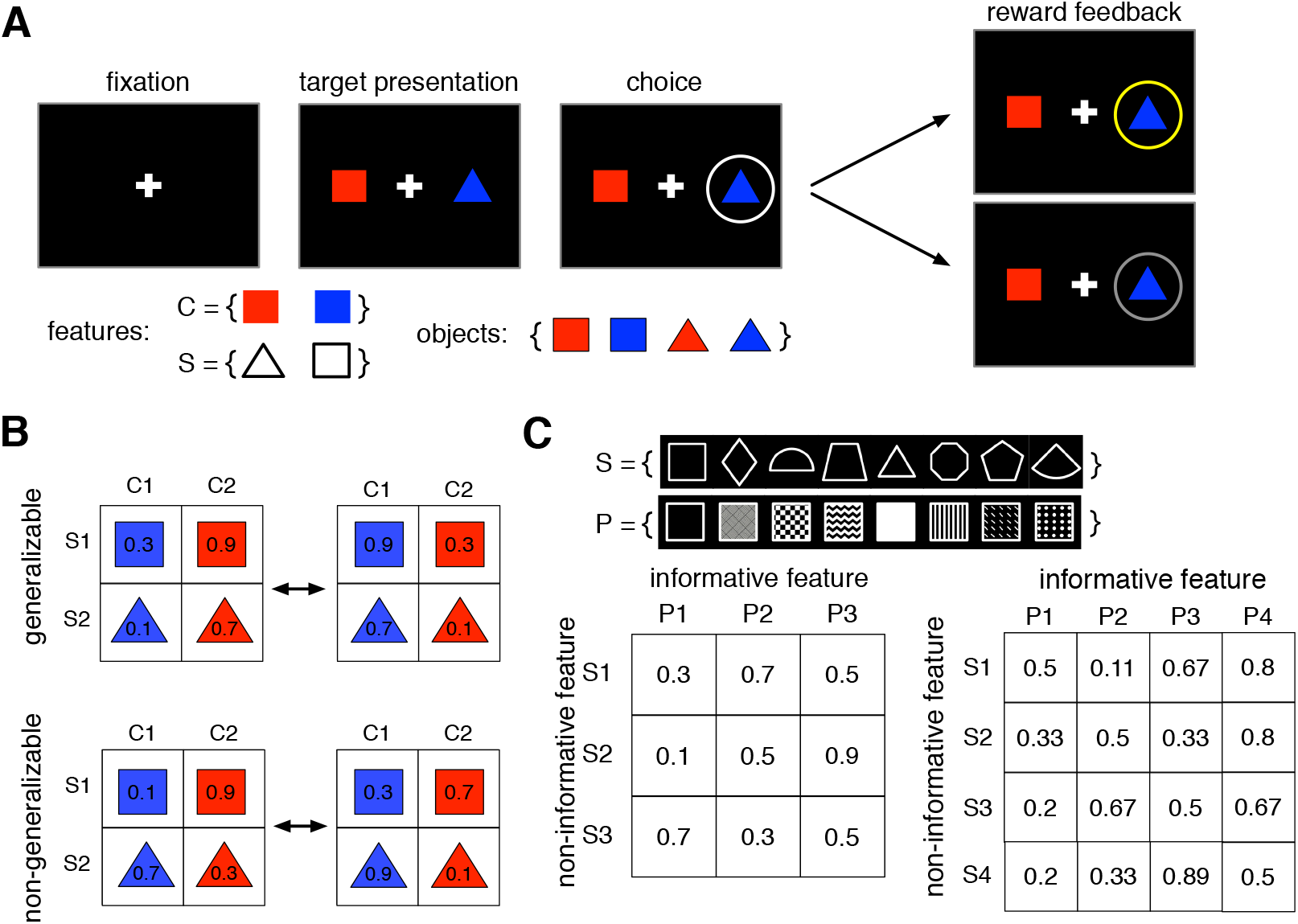
The schematic of the multi-dimensional learning and decision-making task. (**A**) Timeline of a trial in Experiments 1 and 2. On each trial, the subject chose between two objects (colored shapes) and was provided with reward feedback (reward or no reward) on the chosen object. The insets show the set of all features (C, color; S, shape) and objects used in Experiments 1 and 2. (**B**) Examples of the set of reward probabilities (indicated on each object) assigned to the four objects in Experiments 1 and 2. The reward probabilities assigned to the four shapes changed after every 48 trials without any cue to the subject. In the generalizable environment (Experiment 1) reward probabilities assigned to objects were predicted by feature values. In the non-generalizable environment (Experiment 2), there was no generalizable relationship between the reward values of individual objects and their features. **(C)**Reward probabilities were assigned to nine or sixteen possible objects during Experiments 3 and 4, respectively. Objects were defined by combinations of two features (S, shape; P, pattern), each of which could take any of three or four values in Experiments 3 and 4, respectively. The inset shows the set of shapes (top row) and patterns (bottom row) used to construct objects. Reward probabilities were assigned such that no generalizable rule predicted the reward probabilities of all objects based on feature values. In addition to choice trials similar to those in Experiments 1 and 2, Experiments 3 and 4 also included estimation trials where the subject estimated the probability of reward for an individual object by pressing one of ten keys on the keyboard.

### Subjects

Subjects were recruited from the student population at Dartmouth College. The exclusion criterion was similar for all experiments and was defined as performance below chance level (0.5). In total, 59 subjects were recruited (34 females) to perform one or both of Experiments 1 and 2 (33 subjects performed both experiments) in a pseudo-randomized order and on separate days. The chance criterion for Experiments 1 and 2 was equal to performance threshold of 0.5406 (equal to 0.5 plus 2 times s.e.m., based on the average of 608 trials after excluding the first 10 trials of each block). As a result, data from a few subjects (8 out of 51 subjects in Experiment 1 and 19 out of 41 in Experiment 2) whose performance fell below this threshold were excluded from the analysis. An additional subject was excluded from Experiment 2 for submitting the same response throughout the experiment. For Experiment 3, 36 additional subjects were recruited (20 females). The exclusion criterion for this experiment was equal to performance threshold of 0.5447 (equal to 0.5 plus 2 times s.e.m., based on the average of 500 trials after excluding the first 30 trials of each session), resulting in exclusion of 9 subjects. For Experiment 4, 36 more subjects were recruited (22 females). A similar exclusion criterion in this experiment resulted in removal of 11 subjects whose performance fell below 0.5404 (equal to 0.5 plus 2 times s.e.m., based on the average of 612 trials after excluding the first 30 trials of each session). No subject had a history of neurological or psychiatric illness. Subjects were compensated with a combination of money and “t-points,” which are extra-credit points for classes within the Dartmouth College Psychological and Brain Sciences department. After a base rate of $10/hour or 1 t-point/hour, subjects were additionally rewarded based on their performance (e.g., number of reward points), by up to $10/hour.

### Ethics statement

All experimental procedures were approved by the Dartmouth College Institutional Review Board, and informed consent was obtained from all subjects before participating in the experiment.

### Experiments 1 and 2

In these experiments, subjects completed two sessions (each composed of 768 trials and lasting approximately one hour) of a choice task during which they selected between a pair of objects on each trial (Figure 1A). Subjects were asked to choose between objects that differed in color and shape (red triangle, blue triangle, red square, blue square) and to select the object that was more likely to provide a reward in order to maximize the total number of reward points.

During each trial, the selection of an object was rewarded independently of the reward probability of the other object. This reward schedule was fixed for a block of trials (block length, *L* = 48), after which it changed to another reward schedule without any signal to the subject. Sixteen different reward schedules were used; eight schedules consisted of generalizable rules for how combinations of features (color or shape) predicted objects’ reward probabilities, and the other eight lacked generalizable rules for how combinations of features predicted objects’ reward probabilities (see Farashahi et al., 2017 for more details). Therefore, the main difference between Experiments 1 and 2 was that the environments in these experiments were composed of reward schedules with generalizable and non-generalizable rules, respectively. Critically, as the subjects moved between blocks of trials, reward probabilities were reversed in order to induce volatility in reward information throughout the task. On each block of the generalizable environment (Experiment 1), feature-based rules could be used to correctly predict reward throughout the task. In the non-generalizable environment (Experiment 2), however, the reward probability assigned to each object could not be determined based on the objects’ features (e.g. in a multiplicative fashion). More details about the reward schedules in Experiments 1 and 2 are provided elsewhere (Farashahi et al., 2017).

### Experiment 3

In this experiment, subjects completed two sessions, each of which included 280 choice trials interleaved with five or eight short blocks of estimation trials (each estimation block with eight trials). Subjects were asked to choose the more rewarding of two objects during each trial of the choice task. These objects were drawn from a set of eight objects constructed using combinations of three distinct patterns and three distinct shapes (Figure 1C). The two objects presented during each trial always differed in both pattern and shape. Other aspects of the choice task were similar to those in Experiments 1 and 2, except that reward feedback was given for both objects rather than just the chosen object in order to accelerate learning. Subjects then provided estimates of each object’s reward during estimation trials. Possible values for these estimates ranged from 5% to 95% in 10% increments. All subjects completed five blocks of estimation trials throughout the task (after trials 42, 84, 140, 210, and 280 of the choice task), and some subjects had three additional blocks of estimation trials (after trials 21, 63, and 252) to better assess changes in estimations over time. Each session of the experiment was approximately 45 minutes in length, with a break between sessions. The second session was similar to the first, but with different sets of shapes and patterns.

Selection of a given object was rewarded (independently of the other object on a given trial) based on a reward schedule with a moderate level of generalizability such that only some objects’ reward probabilities could be determined based on their feature values. In Experiment 3, one feature (shape or pattern) was informative about reward probability, while the other feature was not. Although the informative feature was predictive of reward on average, this prediction was not generalizable to all individual objects. Additionally, the non-informative feature did not follow any generalizable rule. That is, the average reward probability across objects with a similar feature value was equal to 0.5 (e.g. shapes S1 to S3 in Figure 1C). This reward schedule ensured that subjects could not use a generalizable feature-based rule to accurately predict reward probability for all objects. Similarly to Experiments 1 and 2, the informative feature was randomly assigned and counter-balanced across subjects to minimize the effects of intrinsic pattern or shape biases. More details about the reward schedules are provided elsewhere (Farashahi et al., 2017).

### Experiment 4

This experiment was similar to Experiment 3, except that each feature could take on one of four values for each feature dimension (i.e. there were four shapes and four patterns), resulting in an environment with higher dimensionality. Each subject completed two sessions of 336 choice trials interleaved with five or eight short blocks of estimation trials (each block with eight trials). The objects in this experiment were drawn from a set of twelve objects, which were combinations of four distinct patterns and four distinct shapes. The probabilities of reward for different objects (the reward matrix) were set such that there was one informative feature and one non-informative feature, just as in Experiment 3 (Figure 1C). Minimum and maximum average reward values for features were similar for Experiments 3 and 4.

### Model fitting

To capture subjects’ learning and choice behavior, we used seven different reinforcement learning (RL) models based on object-based or feature-based learning, which rely on different assumptions about how reward values are represented and updated (see below). These models were fit to experimental data by minimizing the negative log likelihood of the predicted choice probability given different model parameters using the ‘fminsearch’ function in MATLAB (MathWorks, Inc., Natick, MA). Three measures of goodness-of-fit were used to determine the best model to account for the behavior in each experiment: average negative log likelihood (-LL), Akaike information criterion (AIC), and Bayesian information criterion (BIC).

### Object-based RL models

This group of models, based on standard RL, directly estimated reward value of each object based on feedback from previous trials. The uncoupled object-based RL model updated only the reward value of the chosen object using separate learning rates for rewarded and unrewarded trials as follows:

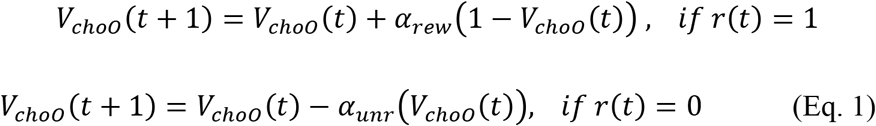

where *t* represents the trial number, *V*_*choO*_ is the estimated reward value of the chosen object, *r*(*t*) is the trial outcome (1 for rewarded, 0 for unrewarded), and *α*_*rew*_ and *α*_*unr*_ are the learning rates for rewarded and unrewarded trials, respectively.

The coupled object-based RL model, on the other hand, updated reward values of both the chosen and unchosen object on each trial in opposite directions. The value of the chosen object was updated with Equation 1, whereas that of the unchosen object was updated as if the reward values were anti-correlated:

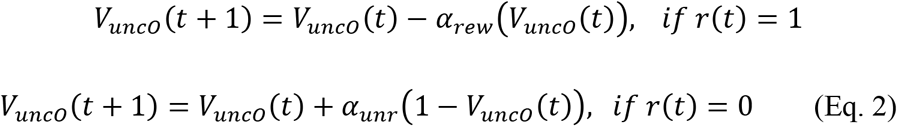

where *V*_*uncO*_ represents the estimated reward value of the unchosen object.

Given the estimated value functions for each object, the probabilities of choosing one of the objects (*O*1 and *O*2) were determined using the following equation:

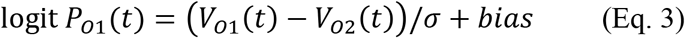

where *P*_*O*1_ is the probability of choosing object 1, *V*_*O*1_ and *V*_*O*2_ are the reward values of the objects presented to the left and right, respectively, *bias* measures a response bias toward the left option in order to capture the subject’s location bias, and *σ* is a parameter measuring the level of stochasticity in the decision process.

### Feature-based RL models

The feature-based RL models utilized the combined reward values of an object’s features to estimate the reward value of an object. Feature reward values were modeled with standard RL, updated similarly to those for the object-based RL model, but where the reward value of the chosen (unchosen) object was instead the chosen (unchosen) feature. For Experiments 1 and 2 in which the alternative objects could have a feature in common (e.g. both were squares), this model resulted in the poorest fit when the reward values of both features of the chosen object were updated. As a result, in the models presented here, we only updated the reward value of the unique feature when there was a shared feature.

Similarly to the object-based RL models, the probability of choosing a shape was modeled with a logistic function:

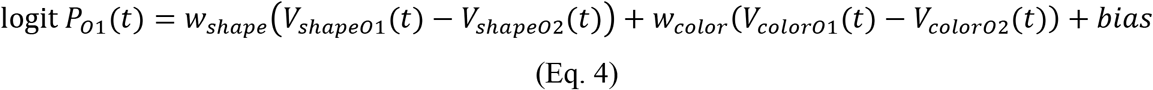

where *V*_*shapeO*1_(*V*_*colorO*1_) and *V*_*shapeO*2_(*V*_*colorO*2_) are the reward values associated with the shape (color) of left and right objects, respectively, *bias* measures a response bias toward the left option, and *w*_*shape*_ and *w*_*color*_ determine the influence of the two features on the final choice.

### RL models with decay

We also modeled the ‘forgetting’ of reward values when objects or features were not chosen. Decay functions, which may capture this phenomenon in multi-dimensional models, were incorporated into the uncoupled models. For unchosen objects or features, reward values decayed to 0.5 (chance or equal information about reward or no reward) with a rate of *d*:

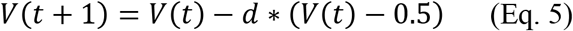

where *t* is the trial number and *V* is estimated reward probability of an object or a feature.

### Analysis of response time

For most analyses, response time (RT) was normalized (z-scored) by first subtracting the mean RT of a given subject from their RT values and then dividing the outcome by the standard deviation of RT for that subject. This normalization allowed us to directly compare data from all subjects. In addition to RT, we also z-scored all independent variables in order to obtain the standardized regression coefficients directly. Standardized regression coefficients quantify the relative influence of predictors measured on different scales. Unlike z-scoring, a constant term in the generalized linear model (GLM) cannot address different ranges of RT for different subjects (variability relative to mean RT), which could result in subjects with more variable RT influencing the results more strongly. Because RTs were z-scored for individual subjects, we did not include an intercept in the GLM. We removed trials with RT less than 50 msec and more than 5 sec since they correspond to choices with no deliberation and when the subject was distracted, respectively.

We used several GLMs with the following regressors (all z-scored) to predict the normalized RT on each trial: 1) the difference between the actual or estimated (using the model that provides the best fit of choice data) reward probabilities of the two objects presented on a given trial; 2) the trial number within a block; 3) reward outcome on the preceding trial; and 4) a ‘model-adoption’ index comparing the prediction of the best feature-based and object-based models on a given trial. The model-adoption index was defined as the difference between the goodness-of-fit by the best feature-based and object-based models using BIC per trial (BIC_p_) defined as:

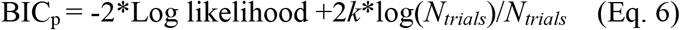

where *k* indicates the number of parameters in a given model. The best object-based and feature-based models were determined based on the BIC for each subject in a given experiment. We note that even though we used BIC for these analyses, similar results were obtained with AIC. Thus the model-adoption index determines the degree to which feature-based or object-based models predicted choice behavior in a given trial more accurately for an individual subject (positive values correspond to a better prediction by the object-based model).

To further investigate the relationship between RT and the adopted model on a given trial, we grouped trials into object-based, feature-based, and equivocal trials depending on which model provided a better fit (based on BIC_p_) and whether the absolute value of model-adoption index was larger than a threshold. More specifically, equivocal trials were defined as trials in which the absolute value of the model-adoption index was less than 0.1. This resulted in assigning approximately 39, 39, and 22 percent of trials as object-based, feature-based, and equivocal, respectively (over all experiments). The percentages of trials assigned to these three types in a given experiment were compatible with the overall learning strategy used in that experiment (Supplementary Table 1).

**Table 1.**
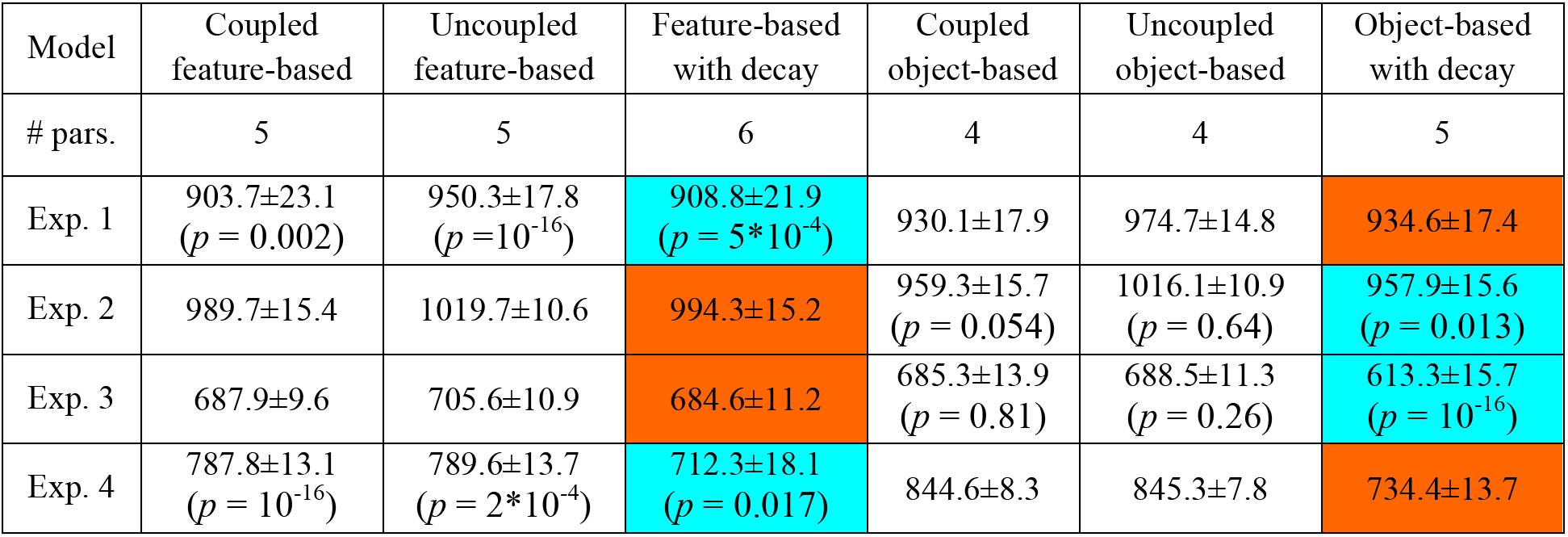
The average values of the Bayesian information criterion (BIC) based on three feature-based RLs and their object-based counterparts, separately for each of four experiments. Reported are the average values of BIC over all subjects (mean±s.e.m.) and *p*-values for comparisons of BIC values between each model and its object-based or feature-based counterparts (two-sided Wilcoxon signed-rank test). The overall best model (feature-based or object-based with decay) and its object-based or feature-based counterpart are highlighted in cyan and brown, respectively.

| Model | Coupled feature-based | Uncoupled feature-based | Feature-based with decay | Coupled object-based | Uncoupled object-based | Object-based with decay |
| --- | --- | --- | --- | --- | --- | --- |
| # pars. | 5 | 5 | 6 | 4 | 4 | 5 |
| Exp. 1 | 903.7±23.1<br>($p = 0.002$) | 950.3±17.8<br>($p = 10^{-16}$) | 908.8±21.9<br>($p = 5*10^{-4}$) | 930.1±17.9 | 974.7±14.8 | 934.6±17.4 |
| Exp. 2 | 989.7±15.4 | 1019.7±10.6 | 994.3±15.2 | 959.3±15.7<br>($p = 0.054$) | 1016.1±10.9<br>($p = 0.64$) | 957.9±15.6<br>($p = 0.013$) |
| Exp. 3 | 687.9±9.6 | 705.6±10.9 | 684.6±11.2 | 685.3±13.9<br>($p = 0.81$) | 688.5±11.3<br>($p = 0.26$) | 613.3±15.7<br>($p = 10^{-16}$) |
| Exp. 4 | 787.8±13.1<br>($p = 10^{-16}$) | 789.6±13.7<br>($p = 2*10^{-4}$) | 712.3±18.1<br>($p = 0.017$) | 844.6±8.3 | 845.3±7.8 | 734.4±13.7 |

We also calculated the relationship between RT and the adopted model in BIC-matched trials. More specifically, for each subject we first picked object-based and feature-based trials for which BICs were between the largest minimum and smallest maximum BIC of object-based and feature-based trials. We then sorted these trials based on their BIC values and computed the mean BIC for each set of trials, and then one by one removed the largest BIC trials from the group with the larger mean BIC, and the smallest BIC trials from the group with the smaller mean BIC. This procedure was repeated until the two means of the BIC values from the two sets of trials were as close to each other as possible. We then calculated the average normalized RT for these two sets of trials.

To test whether there was any effect of previous participation on RT for subjects who performed both Experiments 1 and 2, we also added ‘previous participation’ as a regressor (equal to 0 and 1 if the current experiment was the first and second experiment, respectively) to our GLMs for fitting RT during these experiments.

## Results

To examine how RT depends on the construction and comparison of reward values, we used various reinforcement learning models to determine whether object-based or feature-based learning was adopted by individual subjects in a given experiment. For each model, we computed Bayesian information criterion (BIC) as the measure of goodness-of-fit in order to determine the best overall model to account for the behavior in each experiment (Table 1). Other measures of goodness of fit such as -log likelihood (-LL), and Akaike information criterion (AIC) also revealed similar results as shown in Supplementary Figure 1. The smaller value for each measure indicates a better fit of choice behavior.

This analysis showed that in Experiment 1 (dynamic generalizable environment), the feature-based models provided better fits than the object-based models, indicating that subjects adopted feature-based learning more often in this experiment. In contrast, subjects more frequently adopted object-based learning in Experiment 2 (dynamic non-generalizable environment). In Experiment 3 (static and non-generalizable environment), subjects more frequently adopted object-based learning. An examination of fits and the pattern of estimations over time revealed that in this experiment, subjects adopted feature-based learning first before slowly switching to object-based learning (Farashahi et al., 2017). In contrast, in Experiment 4 (static non-generalizable environment), which involved a larger number of objects and slightly larger generalizability than Experiment 3, subjects more frequently adopted feature-based learning and did not switch to object-based learning.

Altogether, the analysis of subjects’ choice behavior demonstrates that subjects adopted different learning strategies depending on the structure of reward in each environment. As mentioned above, each of these learning strategies requires a different method for the construction and/or comparison of reward value. Object-based learning relies on directly representing and updating the rewards of individual options. In contrast, feature-based learning requires representing and updating the average reward values of all features, and then combining these values to estimate the reward values for individual options. To test whether the learning strategy influenced RT, we next examined the pattern of RT in our four experiments.

The average RTs across all subjects (using median RT of all trials for each subject) were equal to 0.78±0.04s, 0.85±0.05s, 1.04±0.05s, and 1.01±0.03s for Experiments 1 to 4, respectively. A longer RT for Experiments 3 and 4 could be attributed to a larger number of objects in those experiments (8 and 12, respectively, in Experiments 3 and 4, rather than 4 in Experiments 1 and 2). Experiments 1 and 2 were identical except for the way in which reward probabilities were assigned to the four objects, resulting in generalizable and non-generalizable environments. Because of the non-generalizable reward schedules in Experiment 2, this experiment was more difficult to perform than Experiment 1. This was reflected in lower average performance (harvested reward) (Wilcoxon rank-sum test, *p* < 0.001; the average performance for Experiments 1 and 2 were equal to 0.564±0.004 and 0.544±0.003, respectively) and a larger number of subjects whose performance did not differ from chance in Experiment 2 relative to Experiment 1 (Pearson’s chi-square test, *P* = .0014; 8 of 51 in Experiment 1, and 19 of 41 in Experiment 2).

Despite the fact that Experiment 2 was more difficult, the average RT across all subjects in Experiment 2 was not statistically different from that in Experiment 1 (Wilcoxon rank-sum test, *P* = 0.2). This observation can be explained by noting that subjects adopted object-based learning in Experiment 2 more often than in Experiment 1 (74% vs. 38% in Experiments 2 and 1, respectively), and the average RT for subjects who adopted object-based learning was smaller than that for subjects who adopted feature-based learning in both experiments (two-sided Wilcoxon rank-sum test, Experiment 1: *P* = 0.01, Experiment 2: *P* = 0.04). More specifically, the average median RTs for subjects who adopted object-based and feature-based learning in Experiment 1 were equal to 0.71±0.06s and 0.85±0.05s, respectively. The average median RTs for subjects who adopted object-based and feature-based learning in Experiment 2 were equal to 0.79±0.07s and 0.88±0.07s, respectively. These results provide preliminary evidence supporting our hypothesis that construction and comparison of reward value is faster when the object-based strategy is adopted.

Given these results, we next utilized generalized linear models (GLMs) to examine the influence of several factors on the pattern of RT in each experiment. These included the absolute value of the difference in the actual or estimated probability of reward on the two objects presented on a given trial (absolute difference in reward probability), the trial number within a given experiment, reward outcome on the preceding trial, and the model-adoption index (equal to BIC_p_ (Ft) – BIC_p_ (Obj); see Methods). We used normalized RT (z-scored for each subject) in order to analyze data from all subjects together.

The results of the GLM fits revealed several important aspects of RT in our experiments (Table 2). First, subjects became faster over time in all experiments by an average of 300-400ms at the end relative to the beginning of the experiments (Figure 2). Moreover, in Experiments 1 and 2, in which there was a reversal in reward probabilities after every block of 48 trials (dynamic environments), RT increased at the beginning of each block and then decreased as subjects learned the reward probabilities associated with the four objects in each block (Figure 3). This resulted in average of 20ms and 40ms decrease in RT over the course of each block in Experiments 1 and 2, respectively.

**Table 2.** Factors influencing RT. We used two sets of GLMs to predict the normalized RT as a function of the absolute difference in the actual (A) or estimated (B) probability of reward on the two objects presented on a given trial (absolute difference in actual/subjective reward probability), the trial number within a block of the experiment, the difference between BIC per trial (BIC_p_) based on the best feature-based and object-based models (i.e., model-adoption index) for a given subject, and the reward outcome on the preceding trial. Reported values are the normalized regression coefficients (±s.e.m.), *p*-values for each coefficient (two-sided t-test), and adjusted R-squared for each experiment. No interaction term was statistically significant and thus, interactions terms are not reported here.

| Regressor | Abs. difference<br>in actual reward<br>prob. | Trial number | BIC<sub>p</sub> (Ft) – BIC<sub>p</sub> (Obj) | Reward<br>outcome on<br>prev. trial | R<sup>2</sup> |
| --- | --- | --- | --- | --- | --- |
| Exp. 1 | -0.05±0.005<br>($p=10^{-16}$) | -0.15±0.006<br>($p=10^{-16}$) | -0.02±0.005<br>($p=4.7*10^{-6}$) | -0.03±0.006<br>($p=1.9*10^{-6}$) | 0.027 |
| Exp. 2 | -0.07±0.008<br>($p=10^{-16}$) | -0.11±0.008<br>($p=10^{-16}$) | -0.02±0.008<br>($p=0.003$) | -0.04±0.008<br>($p=10^{-16}$) | 0.020 |
| Exp. 3 | -0.06±0.008<br>($p=10^{-12}$) | -0.16±0.008<br>($p=10^{-16}$) | -0.03±0.008<br>($p=0.004$) | -0.05±0.008<br>($p=1.9*10^{-7}$) | 0.031 |
| Exp. 4 | -0.04±0.007<br>($p=10^{-16}$) | -0.23±0.007<br>($p=10^{-16}$) | -0.01±0.007<br>($p=0.01$) | -0.03±0.007<br>($p=7.8*10^{-4}$) | 0.057 |

**Table 2.** Factors influencing RT.
| Regressor | Abs. difference<br>in subjective<br>reward prob. | Trial number | BIC<sub>p</sub> (Ft) – BIC<sub>p</sub> (Obj) | Reward<br>outcome on<br>prev. trial | R<sup>2</sup> |
| --- | --- | --- | --- | --- | --- |
| Exp. 1 | -0.13±0.006<br>($p=10^{-16}$) | -0.16±0.005<br>($p=10^{-16}$) | -0.02±0.005<br>($p=4.4*10^{-4}$) | -0.05±0.006<br>($p=10^{-16}$) | 0.041 |
| Exp. 2 | -0.13±0.008<br>($p=10^{-16}$) | -0.12±0.008<br>($p=10^{-16}$) | -0.03±0.008<br>($p=4.6*10^{-5}$) | -0.07±0.008<br>($p=10^{-16}$) | 0.031 |
| Exp. 3 | -0.21±0.008<br>($p=10^{-16}$) | -0.14±0.008<br>($p=10^{-16}$) | -0.06±0.008<br>($p=0.005$) | -0.04±0.008<br>($p=1.4*10^{-9}$) | 0.073 |
| Exp. 4 | -0.18±0.007<br>($p=10^{-16}$) | -0.21±0.007<br>($p=10^{-16}$) | -0.01±0.007<br>($p=0.03$) | -0.03±0.007<br>($p=1.6*10^{-5}$) | 0.089 |

**Figure 2.**
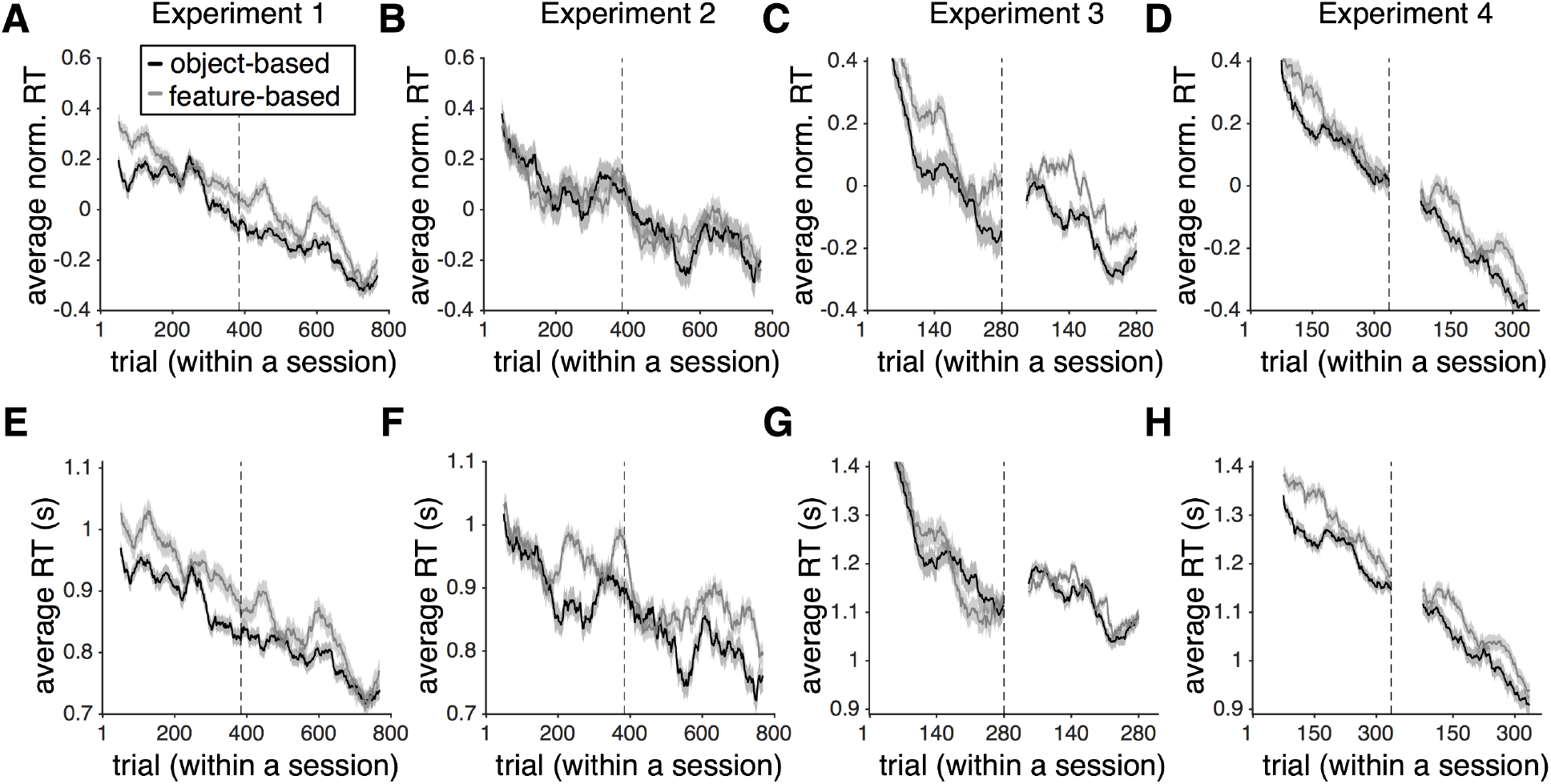
Subjects made decisions more quickly as they progressed through the experiments and during trials when they used the object-based rather than the feature-based model. (**A-D**) Plotted is the average z-scored RT (across all subjects) as a function of the trial number within a session of an experiment separately for trials in which the object-based or feature-based model was used. Panels (A-D) correspond to Experiments 1 to 4, respectively, and shaded areas indicate s.e.m. The gray dotted line shows the time point during a session (in terms of trial number) at which the experiment was paused for a short rest. In Experiments 3 and 4, a different set of objects was introduced at this point. (**E-H**) The same as in (A-D) but showing the average actual RT across all subjects. Overall, subjects made decisions 300-400ms faster at the end relative to the beginning of the experiments. Nevertheless, they were consistently faster on trials when they adopted an object-based rather than feature-based strategy.

**Figure 3.**
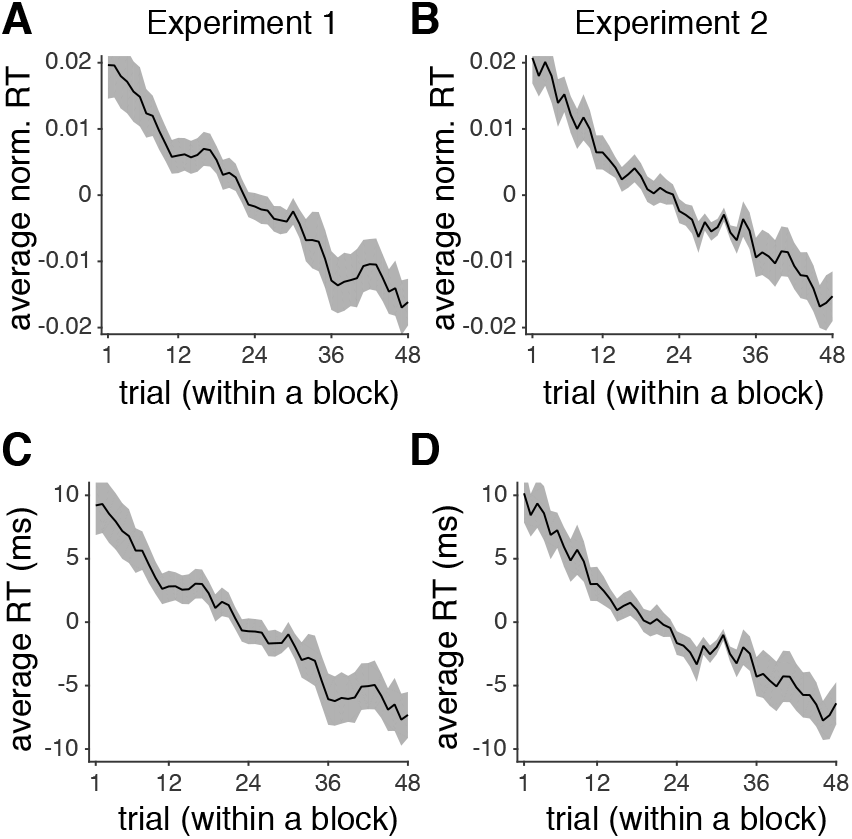
Subjects made decisions more quickly as they progressed through the trials within each block of Experiments 1 and 2. Plotted is the average z-scored RT (across all subjects) as a function of the trial number within a session of Experiment 1 (**A**) and Experiment 2 (**B**). Shaded areas indicate s.e.m. (**C-D**) The same as in (A-B) but showing the average actual RT over all subjects after removing the median RT from each individuals’ RT distribution.

Second, we found that the absolute difference in the actual or estimated reward probabilities negatively affected RT such that RT was maximal when this difference was close to zero (Table 2; Figures 5, 6). Importantly, the absolute difference in estimated reward probabilities (based on the fit of choice behavior of each subject) was a better predictor than the absolute difference in the actual reward probabilities (compare *R*^*2*^ based on the GLMs in panels A and B of Table 2). The absolute difference in the estimated reward probabilities resulted in about 120ms change in RT (Figure 6). Third, the reward outcome on the previous trial also led to shorter RT after rewarded trials compared to after unrewarded trials. Overall, RT on trials following rewarded trials was shorter than RT following unrewarded trials by 33±12ms, 50±24ms, 50±11ms, and 16±13ms in Experiments 1 to 4, respectively, indicating that subjects spent more time making a choice following unrewarded trials.

**Figure 4.**
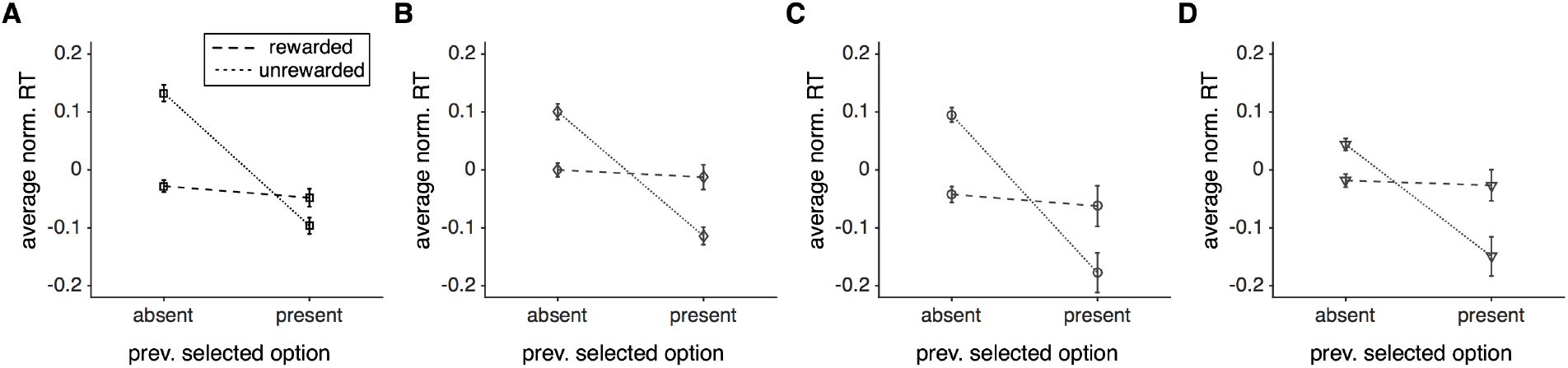
The average (across subjects) normalized RT for different types of trials depending on the presence or absence of previously selected object and rewarded or unrewarded outcome. RT was mainly modulated by the presence or absence of the previously selected object when the previous trial was not rewarded.

**Figure 5.**
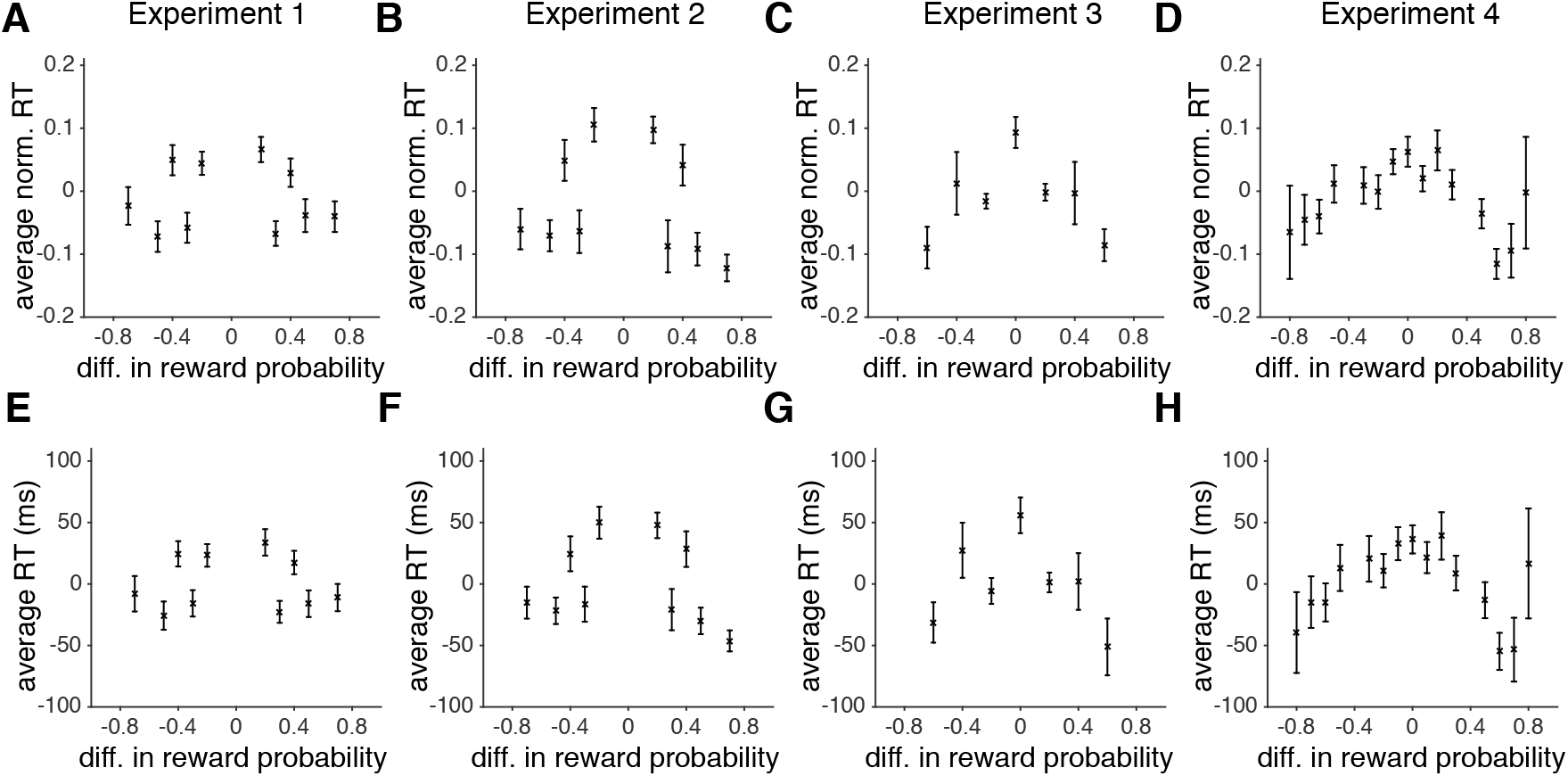
Subjects were faster on trials in which the difference between the actual reward probabilities of the competing objects deviated more from zero. (**A-D**) Panels A-D correspond to Experiments 1 to 4, respectively, and the error bars indicate s.e.m. (**E-H**) The same as in A-D but showing the average actual RT (median removed) over all subjects. Overall, the difference in actual reward probabilities of the two alternative objects resulted in up to ~100 ms increase in RT when reward probabilities were close to each other relative to when they were very different.

**Figure 6.**
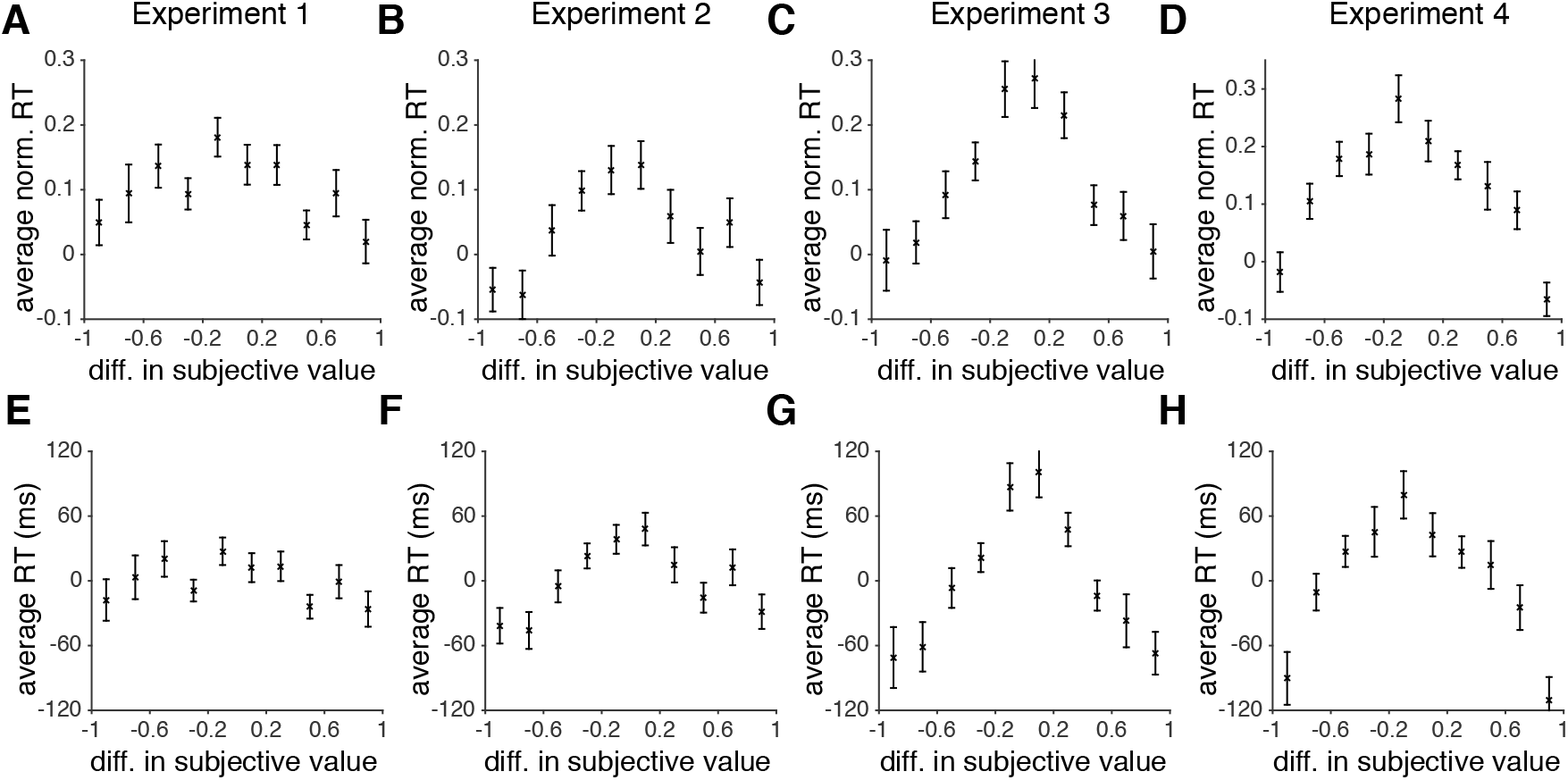
Subjects were faster on trials in which the difference between the estimated reward probabilities of the competing objects deviated more from zero. (**A-D**) Panels A-D correspond to Experiments 1 to 4, respectively, and the error bars indicate s.e.m. (**E-H**) The same as in A-D but showing the average actual RT (median removed) over all subjects. Overall, the difference in estimated reward probabilities of the two alternative objects resulted in up to ~150 ms increase in RT when reward probabilities were close to each other relative to when they were very different.

Previously, it has been shown that the modulation of RT by reward on the preceding trial depends on whether or not the rewarding feature of the target stayed the same between the two trials (Hickey, Chelazzi, & Theeuwes, 2010). Therefore, we also examined whether the effect of reward on the following trial depends on the presence of the previously selected object. To do so, we fit RT data with a new GLM with additional terms: 1) whether the object selected on the previous trial was present or not (previously selected present), and 2) the interactions between reward outcome on the previous trial and previously selected object being present/absent. The results of this GLM not only revealed that presentation of an object selected on the previous trial reduced RT but also showed a strong interaction between this effect and reward outcome on the previous trial (Table 3). Therefore, we next computed the average RT in four sets of trials depending on whether the preceding trial was rewarded or not and whether the option selected on the previous trial was present or absent.

**Table 3.** Predicting RT with a GLM that also includes whether the object selected on the preceding trial was present or absent. We used a GLM to predict the normalized RT as a function of the absolute difference in the estimated (subjective) probability of reward of the two objects presented on a given trial, the trial number within a block of the experiment, the difference between BIC per trial (BIC_p_) based on the best feature-based and object-based models for a given subject, the reward outcome on the previous trial, presence of an object that was selected on the previous trial (prev. selected present), and the interaction of the last two predictors. Reported values are the normalized regression coefficients (±s.e.m.), *p*-values for each coefficient (two-sided t-test), and adjusted R-squared for each experiment.

| Regressor | Abs. difference in subjective reward prob. | Trial number | BIC<sub>p</sub> (Ft) – BIC<sub>p</sub> (Obj) | Reward outcome on prev. trial | Prev. selected present | Reward outcome on prev. trial x prev. selected present | R<sup>2</sup> |
| --- | --- | --- | --- | --- | --- | --- | --- |
| Exp. 1 | -0.12±0.006<br>(*p* = 10<sup>-16</sup>) | -0.16±0.005<br>(*p* = 10<sup>-16</sup>) | -0.02±0.005<br>(*p* = 10<sup>-5</sup>) | -0.05±0.005<br>(*p* = 10<sup>-16</sup>) | -0.05±0.005<br>(*p* = 10<sup>-16</sup>) | -0.05±0.006<br>(*p* = 10<sup>-16</sup>) | 0.045 |
| Exp. 2 | -0.13±0.008<br>(*p* = 10<sup>-16</sup>) | -0.12±0.008<br>(*p* = 10<sup>-16</sup>) | -0.03±0.008<br>(*p* = 10<sup>-5</sup>) | -0.06±0.008<br>(*p* = 10<sup>-16</sup>) | -0.05±0.008<br>(*p* = 4.3*10<sup>-9</sup>) | -0.07±0.008<br>(*p* = 10<sup>-16</sup>) | 0.037 |
| Exp. 3 | -0.21±0.008<br>(*p* = 10<sup>-16</sup>) | -0.13±0.008<br>(*p* = 10<sup>-16</sup>) | -0.06±0.008<br>(*p* = 0.005) | -0.05±0.008<br>(*p* = 10<sup>-9</sup>) | -0.05±0.008<br>(*p* = 6.6*10<sup>-9</sup>) | -0.03±0.008<br>(*p* = 10<sup>-4</sup>) | 0.074 |
| Exp. 4 | -0.18±0.007<br>(*p* = 10<sup>-16</sup>) | -0.20±0.007<br>(*p* = 10<sup>-16</sup>) | -0.01±0.007<br>(*p* = 0.03) | -0.03±0.007<br>(*p* = 10<sup>-5</sup>) | -0.02±0.007<br>(*p* = 2*10<sup>-4</sup>) | -0.02±0.007<br>(*p* = 0.002) | 0.089 |

Interestingly, we found that subjects were faster following rewarded than unrewarded trials when the object presented on the preceding trial was not present, and this was true for all experiments (Figure 4). In contrast, when the previously selected object was present, subjects were faster following unrewarded trials (see Table 7 below for detailed statistics). Overall, there was a larger proportion of trials in which the previously selected object was absent rather than present, resulting in an overall faster RT for rewarded than unrewarded trials. However, subjects were faster when the previously selected object was present rather than absent, but only after unrewarded trials. Therefore, we also found differential effects of previous reward on RT depending on the presence or absence of the object selected on the preceding trial, similarly to previous experiments.

Finally, we found that the model-adoption index (equal to BIC_p_ (Ft) – BIC_p_ (Obj), measuring the degree to which the object-based model provides a better fit than the feature-based model on a given trial) had a negative effect on RT (Table 2). This illustrates that subjects were faster on trials in which the object-based model provided a better fit (i.e., when they adopted object-based learning). To further investigate this relationship, we computed RT for object-based, feature-based, and equivocal trials (trials where the adopted model was ambiguous; see Methods). This analysis illustrated that subjects were faster by about 40ms when they adopted an object-based compared to a feature-based model (Figure 7A-B; Table 3). Moreover, RT was fastest on equivocal trials. At the same time, however, the minimum BIC based on the two best models on each trial, a measure of the best prediction by either model, was the largest for equivocal trials (Figure 7C; Table 4). These indicate that subjects were slightly faster on equivocal trials, perhaps because on those trials they made decisions more randomly as suggested by the reduced quality of fit. To address the concern that faster RT on object-based trials could also be due to more random choice behavior on those trials, we calculated the average normalized RT on feature-based and object-based trials after matching the BIC values (see Methods). We found that even on the BIC-matched trials (i.e. trials with similar quality of fit) subjects were still faster when they adopted the object-based learning compared to when they adopted feature-based learning (one-sided ranksum test, *p* = 0.03, using data from all four experiments; Figure 7D). Therefore, even though the quality of fit was in general worse for object-based trials, faster RT on these trials was not due to more random behavior on those trials. Overall, these results illustrate that RT is influenced by the learning strategy adopted by subjects on a trial-by-trial basis.

**Table 4.** Comparisons of RT and minimum BIC during different types of trials. Reported are the *p*-values (two-sided signed-rank test) for comparisons of the average RT, average normalized RT, and minimum BIC between a given pair of trials (object-based, feature-based, and equivocal trials) across subjects depicted in Figure 7A-C.

|  | Object-based vs.<br>equivocal | Object-based vs.<br>Feature-based | Feature-based<br>vs. equivocal |  |
| --- | --- | --- | --- | --- |
| Exp. 1 | 0.0023 | 0.0039 | 0.0001 | average RT<br>(ms) |
| Exp. 2 | 0.0024 | 0.2305 | 0.0051 |  |
| Exp. 3 | 0.6139 | 0.0046 | 0.0003 |  |
| Exp. 4 | 0.0578 | 0.0480 | 0.0009 |  |
| Exp. 1 | 0.0045 | 0.0030 | 0.0001 | average<br>norm. RT |
| Exp. 2 | 0.0026 | 0.2443 | 0.0064 |  |
| Exp. 3 | 0.5806 | 0.0062 | 0.0003 |  |
| Exp. 4 | 0.0653 | 0.0544 | 0.0010 |  |
| Exp. 1 | 0.0036 | 0.0001 | 0.1231 | minimum<br>BIC |
| Exp. 2 | 0.0001 | 0.0001 | 0.0765 |  |
| Exp. 3 | 0.5615 | 0.0001 | 0.0001 |  |
| Exp. 4 | 0.0002 | 0.0001 | 0.0032 |  |

**Figure 7.**
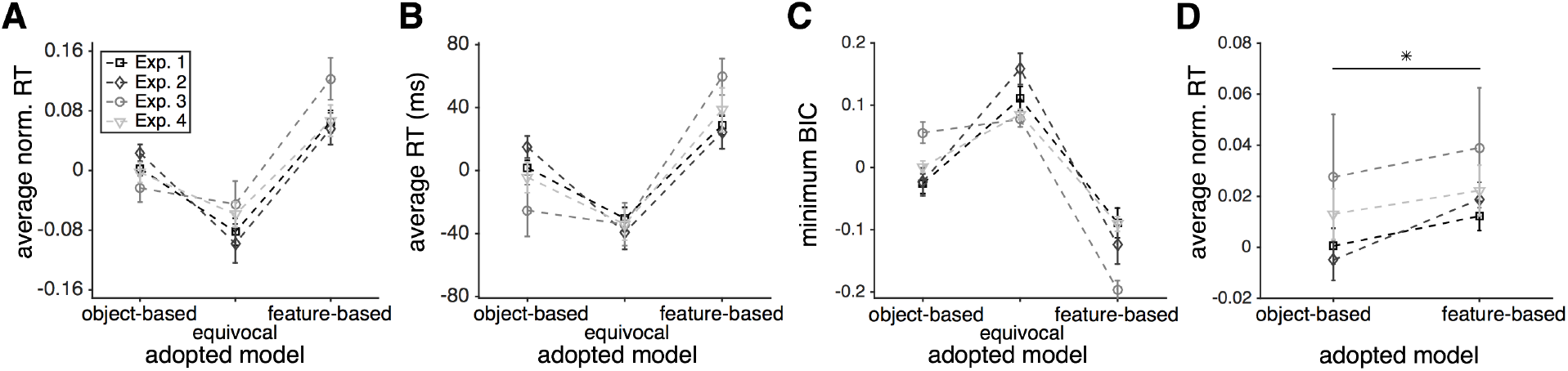
The RT depended on the model adopted by individual subjects on a trial-by-trial basis. (**A**) Plotted is the average (across subjects) normalized RT for three types of trials. The error bars indicate s.e.m. Subjects made decisions faster during trials where the object-based model was used than when the feature-based model was used. In some experiments, subjects were even faster on equivocal trials (when the better model was ambiguous). (**B**) The same as in (A) but showing the average actual RT over all subjects after removing the median RT from each individual RT distribution. **(C)** Plotted is the average (across subjects) of minimum BIC for each group of trials after removing the average minimum BIC over all trials for each subject. Subjects’ choice behavior was hardest to fit on equivocal trials indicating that they made choices more stochastically on those trials. (**D**) Plotted is the average (across subjects) normalized RT on BIC-matched object-based and feature-based trials. Similarly to the result obtained with all trials, on BIC-matched trials, subjects made decisions faster during trials when they adopted the object-based rather than feature-based strategy.

Because subjects also became faster over time, a shorter RT on trials when the object-based strategy was adopted could be exaggerated by a gradual shift from using feature-based to object-based strategy and shorter RT as each experiment progressed (especially in Experiment 3 in which this shift was prominent). To test this possibility, we also fit RT using a GLM model that included all interaction terms between the four predictors in Table 2. None of these interactions and especially the interaction between trial number and model-adoption index (measured by the difference in BIC_p_ values) were statistically significant (*p* > .05). We also calculated the average RT over the course of the experiment across all subjects separately for trials in which object-based or feature-based strategy was adopted (Figure 2). We found a consistently shorter RT on object-based trials during the entire course of each experiment. Taken together, these results suggest that the overall faster RT on object-based trials was not caused by a combination of a shift to object-based learning and faster decision making as subjects progressed through each experiment.

Because a large number of subjects performed in both Experiments 1 and 2, we also tested the effect of previous participation on RT in a given experiment (see Methods). This analysis did not reveal any effects of previous participation on RT. We also used a GLM to predict the average value of the difference in the goodness-of-fit (<BIC_p_(Ft) – BIC_p_(Obj)>) based on the current experiment (1 or 2), previous participation, and their interactions (capturing the order of experiments). The result of this GLM did not reveal any effect of previous participation or the order of experiments on the learning strategy.

Finally, to ensure that exclusion criterion did not bias our results in a certain way, we analyzed RT of excluded subjects using the same models and methods used for included subjects. We found higher BIC values for excluded compared to included subjects, suggesting that on average, choice behavior was more difficult to fit for removed subjects (Supplementary Table 3). More importantly, we found that the overall best model based on object-based or feature-based strategy (models with decay) did not provide a significantly better fit than the counterpart model for excluded subjects. This indicates that the adopted model could not be detected well in excluded subjects. Nevertheless, we also analyzed RT data from excluded subjects in all four experiments using the same GLM used for included subjects (Supplementary Table 4). This analysis revealed that the dependence of RT on the four regressors was similar for included and excluded subjects during Experiments 2 and 4 (experiments with largest percentages of excluded subjects), although the effect of the difference in BIC_p_ was not significant for Experiment 4. The signs of all other regressors with a significant effect on RT were similar for excluded and included subjects. Altogether, these results demonstrate that excluded subjects likely did not engage in the experiment or use any certain strategy. More importantly, our exclusion criterion did not lead to removal of meaningful information nor did it bias our conclusion.

## Discussion

We investigated the pattern of RT during a set of a multi-dimensional learning and decision-making tasks in order to provide insights into neural mechanisms underlying the construction and comparison of reward values. Specifically, we tested whether the type of learning strategies adopted by the subjects and the underlying reward value representation were reflected in the pattern of RT during binary choice. We found a decrease in RT over the course of experiments and after rewarded trials. Moreover, RT was strongly modulated by the difference in the value of reward probabilities assigned to the two alternative objects on a given trial such that it was maximal when the absolute value of the difference in the actual or estimated reward probabilities was minimal. As expected, this effect was stronger for estimated than for actual reward probabilities. More importantly, the model adopted by the subject on a trial-by-trial basis influenced RT such that subjects were faster when they made a choice using the object-based rather than the feature-based strategy. These results confirm our hypotheses that at least two stages of value-based decision making, construction of reward value for each option and value comparison between pairs of options, are influenced by the learning strategy adopted by the subject. A crucial aspect of our experiments, which allowed us to study the influence of value construction on RT, is the rich multi-dimensional task in which multiple learning strategies coexist and interact to determine choice.

The subjects in our experiments could perform the task as a search (i.e. foraging) for more rewarding objects or features on each trial (especially during Experiments 3 and 4 with a large set of objects). Therefore, our results could be relevant to visual search and object identification based on reward values (Wolfe, 2013). We found that RT for selection between two options increased as the subjective values of the two options became more similar. If we consider selection between the two objects on a given trial as a search for a more rewarding object (target), our observation dovetails with the finding in visual search that more visual similarity between the target and distractor increases RT (Duncan & Humphreys, 1989). Interestingly, there are a number of studies in which the visual search has been manipulated by differential reward assignment on targets or distractors (Anderson, Laurent, & Yantis, 2011; Hickey et al., 2010; Hickey & Peelen, 2015; Theeuwes & Belopolsky, 2012). Overall, those studies have demonstrated that task-irrelevant distractors previously associated with larger rewards can slow down target detection (Anderson et al., 2011; Hickey & Peelen, 2015), and this reduction in performance has been attributed to attentional capture by the rewarding feature. Interestingly, in an experiment in which different reward magnitudes (large and small) were randomly delivered on correct trials, the magnitude of reward differentially modulated RT on the following trial depending on whether the rewarding feature of the target stayed the same or not between the two trials (Hickey et al., 2010). If the feature stayed the same, subjects were faster after high-reward trials relative to low-reward trials. In contrast, if the feature switched, subjects were slower after high-reward trials relative to low-reward trials.

Our results also could be linked to the distance effect in numerical cognition where the reaction time for discriminating two numbers decreases as the difference between the two numbers increases (Moyer & Bayer, 1976; Moyer & Landauer, 1967). Interestingly, the distance effect seems to be influenced more strongly by response-related processes than overlap between the representations of numbers (Van Opstal, Gevers, De Moor, & Verguts, 2008). The response time in our experiments, however, is influenced by both the difference between the estimated values of competing objects on each trial (value comparison) and how their values are constructed (representation of value).

The feature-based learning provides a heuristic for learning and choice in dynamic, multi-dimensional environments. Importantly, heuristic feature-based learning not only reduces dimensionality but also increases adaptability without compromising precision (Farashahi et al., 2017). Here, we showed that subjects were faster when they made a choice using the object-based rather than the feature-based model. Based on these results, we predict that there will be a larger difference in RT when feature-based versus object-based strategy is adopted for more high-dimensional options. However, RT using the feature-based strategy could be dramatically reduced for high-dimensional options if the comparison between the two options is limited to dissimilar features only, allowing the feature-based learning to be a “fast and frugal” heuristic (Gigerenzer & Goldstein, 1996). The dependence of RT on the subject’s adopted model of the environment also implies that the adopted model should be estimated and considered when interpreting the top-down effects during conventional or foraging visual search tasks (Wolfe, 2013; see Khorsand, Moore, & Soltani, 2015 for a review of top-down effects).

Finally, with regard to how value comparison contributes to RT during decision making, we found that the difference in the probability of reward assigned to the two targets presented on each trial predicted RT, similarly to our previous observation (Soltani, De Martino, & Camerer, 2012). There is growing interest in economics to use RT in order to infer a decision maker’s preference beyond what can be revealed only by choice (Clithero, 2016a, 2016b; Konovalov & Krajbich, 2016; Spiliopoulos & Ortmann, 2016). For example, based on the observation that RT increases as the subjective values of options become closer to each other in a binary choice, Konovalov and Krajbich (Konovalov & Krajbich, 2016) have suggested that RT could be used to detect the indifference point. In another work, Clithero (Clithero, 2016a) has suggested that RT can be used along with choice to improve out-of-sample predictions. Our results suggest that even the adopted model could be predicted based on RT on a given trial. It would be interesting to test in the future whether RT can be used to improve the fit of choice behavior. However, as we showed here, RT monotonically decreases over time even during continuous learning, indicating that precautions must be taken if RT is used as a measure of the indifference point in such situations.

Altogether, by examining behavior during a set of complex, multi-dimensional learning and decision-making tasks, we show that the pattern of RT can be used to investigate different stages of value-based leaning and decision making, and more specifically, how reward value is constructed and compared on a single trial.

## Acknowledgments

This work was supported by NSF EPSCoR (RII Track-2 FEC) grant to A.S.

---

**Supplementary Figure S1.**
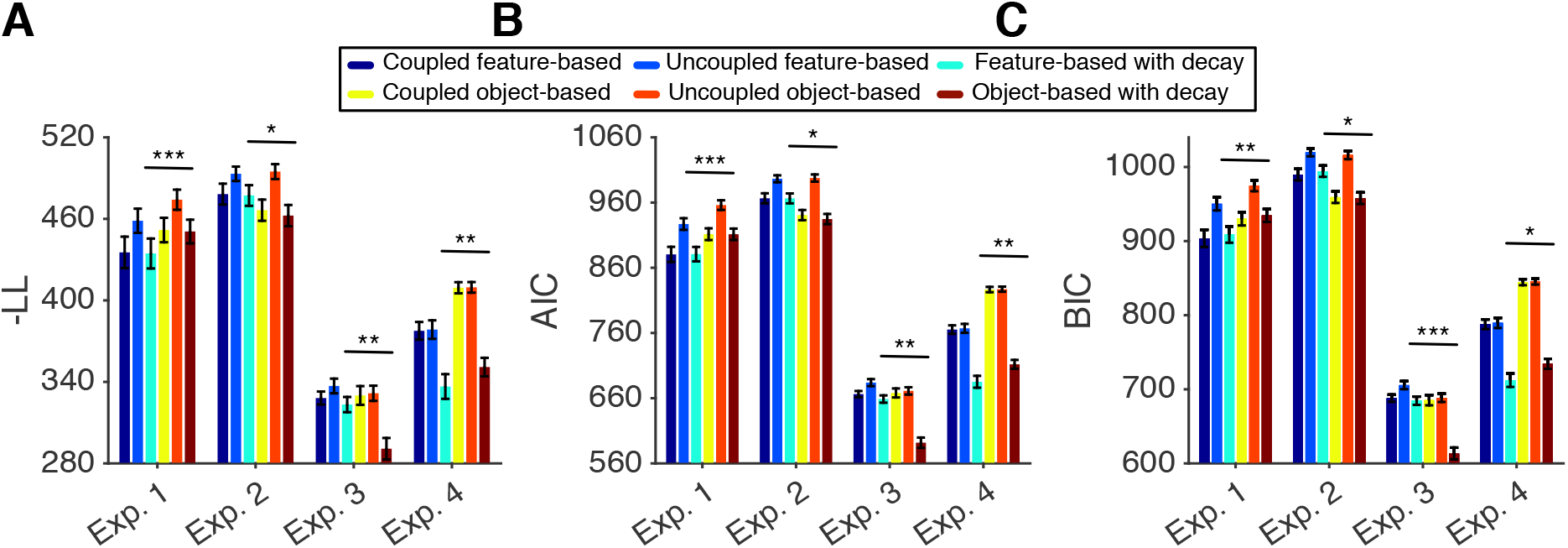
Comparison of the goodness-of-fit measures in all experiments. Panels (A-C) plot the goodness-of-fit measures in terms of the, negative log likelihood (-LL), Akaike information criterion (AIC), and Bayesian information criterion (BIC), respectively. The goodness-of-fit values are computed by averaging over all subjects (mean ± s.e.m.) separately for three feature-based RLs and their object-based counterparts and for Experiments 1 to 4. The significance level of the comparison between each model that provides the best fit in a given experiment and its object-based or feature-based counterpart is coded as: 0.01 < *P* < 0.05 (*), 0.001 < *P* < 0.01 (**), and *P* < 0.001 (***).

**Supplementary Table 1.** Percentage of subjects who adopted feature-based learning and percentages of trials grouped as feature-based, equivocal, and object-based in each experiment. Reported values are mean±std for each experiment. During Experiments 1 and 4, more subjects adopted feature-based learning whereas during Experiments 2 and 3 more subjects adopted object-based learning. The differences between the percentages of trials identified as feature-based and object-based were compatible with the overall adopted learning strategy in each experiment.

| Experiment<br>(no. subjects) | % Feature-based<br>subjects (# subjects) | % Feature-based<br>trial | % Equivocal<br>trials | % Object-based<br>trials |
| --- | --- | --- | --- | --- |
| Exp. 1 (n = 43) | 88 (n = 38) | 40.2±9.2 | 27.9±12.3 | 31.9±7.2 |
| Exp. 2 (n = 21) | 24 (n = 5) | 38.3±8.7 | 18.0±7.3 | 43.7±10.4 |
| Exp. 3 (n = 27) | 30 (n = 8) | 32.7±7.5 | 17.1±5.8 | 50.1±11.9 |
| Exp. 4 (n = 25) | 76 (n = 19) | 43.1±6.9 | 23.3±3.3 | 33.1±7.4 |

**Supplementary Table 2.** Comparisons of RT during different types of trials depending on the presence or absence of previously selected object and rewarded or unrewarded outcome. Reported are the *p*-values (two-sided signed-rank test) for comparisons of the average RT between a given pair of trials (across subjects) depicted in Figure 4.

|  | abs&rew vs.<br>abs&unrew | pres&rew vs.<br>pres&unrew | pres&rew vs.<br>abs&rew | abs&unrew vs.<br>pres&unrew | pres&rew vs.<br>abs&unrew | abs&rew vs.<br>pres&unrew |
| --- | --- | --- | --- | --- | --- | --- |
| Exp. 1 | $5.8 \times 10^{-7}$ | 0.0425 | 0.4688 | $3.4 \times 10^{-8}$ | $1.8 \times 10^{-5}$ | 0.0490 |
| Exp. 2 | 0.0078 | 0.0355 | 0.9861 | $6.9 \times 10^{-5}$ | 0.0355 | 0.0046 |
| Exp. 3 | $2.6 \times 10^{-5}$ | 0.0088 | 0.3488 | $3.5 \times 10^{-5}$ | $1.0 \times 10^{-3}$ | 0.0023 |
| Exp. 4 | 0.0049 | 0.0087 | 0.6377 | $3.0 \times 10^{-4}$ | 0.1155 | 0.0074 |

**Supplementary Table 3.** The average values of the Bayesian information criterion (BIC) based on three feature-based RLs and their object-based counterparts, separately for excluded subjects in each of four experiments. Reported are the average values of BIC over all subjects (mean±s.e.m.) and *p*-values for comparisons of BIC values between each model and its object-based or feature-based counterparts (two-sided Wilcoxon signed-rank test). The overall best model (feature-based or object-based with decay) and its object-based or feature-based counterpart are highlighted in cyan and brown, respectively.

| Model | Coupled feature-based | Uncoupled feature-based | Feature-based with decay | Coupled object-based | Uncoupled object-based | Object-based with decay |
| --- | --- | --- | --- | --- | --- | --- |
| # pars. | 5 | 5 | 6 | 4 | 4 | 5 |
| Exp. 1 | 1008.4±22.4<br>(*p* = 0.43) | 1016.9±22.9<br>(*p* = 0.82) | 1014.6±23.0<br>(*p* = 0.49) | 1017.1±20.2 | 1036.1±21.0 | 1025.4±20.3 |
| Exp. 2 | 994.9±27.4 | 1026.5±25.3 | 993.8±28.1 | 989.2±25.3<br>(*p* = 0.62) | 1015.7±24.6<br>(*p* = 0.03) | 987.4±24.8<br>(*p* = 0.40) |
| Exp. 3 | 746.5±10.2 | 734.8±12.0 | 698.2±11.8 | 777.2±5.9<br>(*p* = 0.03) | 750.1±11.6<br>(*p* = 0.20) | 698.3±13.0<br>(*p* = 0.57) |
| Exp. 4 | 901.6±11.5<br>(*p* = 0.003) | 870.7±24.8<br>(*p* = 0.042) | 840.1±31.4<br>(*p* = 0.68) | 924.9±5.8 | 905.2±11.3 | 871.3±25.2 |

**Supplementary Table 4.** Factors influencing RT for excluded subjects. We used a GLM to predict the normalized RT as a function of the estimated probability of reward of the two objects presented on a given trial (absolute difference in subjective reward probability), the trial number within a block of the experiment, the difference between BIC per trial (BIC_p_) based on the best feature-based and object-based models (i.e., model-adoption index) for a given subject, and the reward outcome on the preceding trial. Reported values are the normalized regression coefficients (±s.e.m.), *p*-values for each coefficient (two-sided t-test), and adjusted R-squared for each experiment. No interaction term was statistically significant and thus, interactions terms are not reported here.

| Regressor | Abs. difference<br>in subjective<br>reward prob. | Trial number | BIC<sub>p</sub> (Ft) – BIC<sub>p</sub> (Obj) | Reward<br>outcome on<br>prev. trial | R<sup>2</sup> |
| --- | --- | --- | --- | --- | --- |
| Exp. 1 | -0.06±0.010<br>($p = 1.4*10^{-9}$) | -0.16±0.010<br>($p = 10^{-16}$) | 0.01±0.009<br>($p = 0.59$) | -0.04±0.010<br>($p = 1.7*10^{-6}$) | 0.030 |
| Exp. 2 | -0.02±0.008<br>($p = 3.5*10^{-11}$) | -0.10 ±0.008<br>($p = 10^{-16}$) | -0.02±0.008<br>($p = 0.004$) | -0.04±0.008<br>($p = 4.7*10^{-7}$) | 0.012 |
| Exp. 3 | -0.12±0.014<br>($p = 10^{-16}$) | -0.10±0.014<br>($p = 2.7*10^{-12}$) | 0.01±0.014<br>($p = 10^{-16}$) | -0.06±0.014<br>($p = 1.8*10^{-6}$) | 0.028 |
| Exp. 4 | -0.04±0.007<br>($p = 10^{-5}$) | -0.24±0.007<br>($p = 10^{-16}$) | -0.01±0.007<br>($p = 0.97$) | -0.05±0.007<br>($p = 3.5*10^{-6}$) | 0.062 |

## Notes

**Conflict of interest:** The authors declare no competing interests.

